# The evolution of host resistance to a virus is determined by resources, historical contingency, and time scale

**DOI:** 10.1101/2022.09.08.507185

**Authors:** Elisa Visher, Hannah Mahjoub, Khadija Soufi, Nilbert Pascual, Vivian Hoang, Lewis J. Bartlett, Katherine Roberts, Sean Meaden, Mike Boots

## Abstract

Hosts can often evolve resistance to parasites (and other stressors), but such resistance is generally thought to be constrained by trade-offs with other traits. These trade-offs determine the host’s optimal resistance strategy and whether resistance cycles, diversifies, and/or is maintained in the absence of parasite. However, trade-offs are often inconsistently measured across experiments and can depend on environmental conditions. Here, we extend a selection experiment evolving resistance to viral infection under variable resource quality in the *Plodia interpunctella* model system to explore the evolutionary conditions leading to an incongruent earlier measurement of costless resistance. We find that environmental resource quality, historical contingency, and the time scale of selection all affect trade-offs in our long-term selection experiment. Specifically, populations selected for resistance with the dual stressor of low resource quality are slowed, but not prevented, from evolving resistance. Second, variation in starting populations or early sampled adaptations led to contingency towards context-dependent resistance. Finally, some costs to resistance observed at early time points were compensated over longer evolutionary time scales. Our work therefore informs perspectives for the predictability of adaptation and how variation in specific evolutionary conditions can alter the evolutionary trajectories of a population towards costly or costless resistance strategies.

## Introduction

Trade-offs between life-history characteristics are critical to evolutionary outcomes and are central to many of our theories for adaptation and diversification (Ackermann and Doebeli, 2004; Agrawal and Lively, 2002; Darwin, 1859; Garland et al., 2022; Levins, 1968). In the case of a host evolving resistance to a pathogen (or other stressor), trade-offs between resistance and other fitness-relevant traits can lead to the evolution of: (1) intermediate (optimal) resistance strategies, (2) diversification through negative frequency dependent selection from ecological feedbacks, (3) resistance cycling, and/or (4) the loss of resistance in the absence of the stressor (Andersson and Hughes, 2010; Best et al., 2010; Boots et al., 2014, 2009; Boots and Haraguchi, 1999; Cotter et al., 2004; Fuxa and Richter, 1989; Graham et al., 2005; Moret and Schmid-Hempel, 2000). More broadly, understanding the costs of resistance is essential to predicting when resistance has a high potential to evolve and persist in scenarios ranging from antibiotic resistance of pathogens (Herren and Baym, 2022), resistance management in the control of invasive species and pests (Kerr et al., 2017), and coevolutionary arms races (Brodie et al., 2002). There are therefore both fundamental and applied reasons to measure both the strength and shape of trade-off relationships (de Mazancourt and Dieckmann, 2004; Ehrlich et al., 2020; Farahpour et al., 2018; Hoyle et al., 2008; Jessup and Bohannan, 2008; Kamo et al., 2007; Kasada et al., 2014; Maharjan et al., 2013; Mealor and Boots, 2006).

Despite the central importance of trade-offs, evolutionary biology has consistently been plagued by the issue that trade-offs are difficult to measure and inconsistently observed (Bono et al., 2017; Cressler et al., 2015; Fry, 1996; Visher and Boots, 2020). This has led to a rich body of empirical work that attempts to discern factors that influence when trade-offs are observed (ex. Bono et al., 2017; Fry, 2003; Stearns, 1989). When trade-offs are not observed, this can sometimes be due to measurement error where costs to some adaptive phenotype exist in fitness dimensions that are not measured (Kawecki, 2020; Kinsler et al., 2020). In other cases, high environmental quality may obscure trade-offs as the organism can allocate resources to buffer costs (Jessup and Bohannan, 2008; Kraaijeveld and Godfray, 1997; Luong and Polak, 2007; McKean et al., 2008). In yet other cases, the likelihood of a population evolving costly (or costless) strategies may be influenced by its specific evolutionary conditions including the environment that it is adapting to, the time scale of selection, the heterogeneity (spatial or temporal) of the environment, and the starting genotypes in the population (Bono et al., 2017; Card et al., 2019; Remold et al., 2008). A better understanding of how these processes define both the nature and our potential to measure trade-offs requires the detailed analysis of well described trade-off relationships.

One of the better characterized trade-offs in evolutionary ecology has been the trade-off between resistance to viral infection and development time in the *Plodia interpunctella* (Indian Meal Moth) (Hübner) and *Plodia interpunctella* granulosis virus (PiGV) model system (Boots and Begon, 1993). The trade-off that increased resistance comes at a cost of longer development time has been established repeatedly by laboratory experimental evolution selecting for resistance (Boots, 2011; Boots and Begon, 1993), by assaying natural populations with phenotypic variation (Boots and Begon, 1995), and by assaying inbred genotypes with phenotypic variation in the absence of infection (Bartlett et al., 2018). It has also been established that costs to resistance can be mediated by the quality of resources provided to the population during evolution so that populations evolving with low-quality resources evolve lower, more costly resistance (Boots, 2011). However, this trade-off has recently proven breakable because laboratory experimental evolution selecting on development time (rather than resistance) results in fast development selected populations actually having higher resistance than their slow development selected counterparts (Bartlett et al., 2020). Additionally, a second laboratory evolution experiment re-evolved resistance under high and low resource quality, as in (Boots, 2011), to explore the genetic basis of resistance (Roberts et al., 2020). In this experiment, the authors saw inconsistent, context-dependent resistance evolution and did not find trade-offs between resistance and growth rate in virus-selected populations evolved in high-quality resources. Furthermore, virus-selected populations evolved in low-quality resources did not significantly evolve resistance and what resistance they do evolve only showed trade-offs in the low-quality, but not common garden, environment. These results therefore provide an exciting opportunity to explore how variation in specific ecological and evolutionary conditions can alter the evolutionary trajectories of a population towards costly or costless resistance strategies.

Compared to (Boots, 2011), (Roberts et al., 2020) use the same selection conditions of constant PiGV-exposure and different resource qualities, but, in contrast, they selected from a genetically distinct starting population. The variation in evolution outcomes may be explained by this difference in starting population genetics. However, the fact that the populations in (Roberts et al., 2020) inconsistently evolve resistance also suggests that the lack of trade-offs could be because these populations are evolutionarily ‘behind’ (despite the (Roberts et al., 2020) lines being assayed after 14 generations of selection, compared to the 10 in (Boots, 2011)) and not yet running into evolutionary costs. In this paper, we examine the repeatability of trade-off observation by testing the hypothesis that differences in evolved resistance and its costs are simply due to slower evolution in the second experiment (Roberts et al., 2020). To do this, we extend the time scale of selection on a subset of the (Roberts et al., 2020) populations and re-assay resistance and development time in common garden and home quality environments. Extending the time scale of selection also allows us to test additional questions about the temporal dynamics of costs and whether they can be compensated over longer time scales (Andersson and Hughes, 2010) and about the longer-term dynamics of evolution under dual stressors (in this case virus infection and low resource quality) (Hiltunen et al., 2018). We therefore explore whether: 1) resistance will evolve if given more time under selection, 2) if the lack of trade-offs in these populations can be explained by them being ‘behind’ in evolution and therefore not at Pareto fronts where resistance phenotypes are constrained (Li et al., 2019; Shoval et al., 2012), and 3) whether longer evolutionary time scales will allow populations evolving under low resource quality to ‘catch-up’ to those evolving in high resources. By exploring these questions, we gain insight into the historical contingency of resistance evolution and its trade-offs.

## Methods

### Study system

*Plodia interpunctella* (Hübner), the Indian meal moth, is a stored grain pest with cyclical population dynamics in the lab. During their five larval instar stages, *P. interpunctella* live at high population densities within the food that they were laid into and consume. After the fifth instar, *P. interpunctella* pupate and eclose. *P. Interpunctella* adult moths do not eat and primarily disperse, mate, and reproduce (Gage, 1995; Mohandass et al., 2007; Silhacek and Miller, 1972). Plodia interpunctella granulosis virus (PiGV) is an obligately lethal, dsDNA baculovirus (Harrison et al., 2016) that infects *P. interpunctella*. Larvae orally ingest PiGV occlusion bodies that have been released into environment or during the process of larval cadaver cannibalism. For successful infection to occur, the virus must shed its protein coat in the gut and infect gut epithelial cells in the budded virus form, cross the gut membrane to establish systemic infection though the larval fat body, and then package into the infectious, protein-coated occlusion body form, at which point the infection kills the larvae. Infection can be cleared before the host is fully infected through various resistance mechanisms including gut ecdysis during molting stages, freeing the host to carry out its natural life cycle and pupate into an adult moth (Boots and Begon, 1993; Engelhard and Volkman, 1995).

### Host line selection and maintenance

Populations of *P. interpunctella* were initially established from an outcrossed *P. interpunctella* population by (Roberts et al., 2020) and selected under 4 treatment conditions: high-quality food with virus (VHF), low-quality food with virus (VLF), high-quality food without virus (CHF), and low-quality food without virus (CLF). Food quality was manipulated by replacing a portion of the cereal mix (50% Ready Brek ©, 30% wheat bran, and 20% ground rice by weight) with either 10% (high-quality food) or 55% (low-quality food) methyl cellulose, a non-digestible fibrous bulking agent, by weight. This alters the amount of nutrition available to the larvae without altering feeding rates (Boots, 2011; Boots and Begon, 1994; Boots and Roberts, 2012). These dry cereal mixtures were then mixed with brewer’s yeast (100g per 500g dry mix), sorbic acid (2.2g per 500g dry mix), methyl paraben (2.2g per 500g dry mix), honey (125mL per 500g dry mix), and glycerol (125mL per 500g dry mix) to form the control (CHF and CLF) food types. For virus food types (VHF and VLF), virus from a stock solution of PiGV was also mixed into the food at a dose corresponding to LD20 for the ancestral *P. interpunctella* population (see (Roberts et al., 2020)).

For our experiment, we maintained 5 replicate selection lines per treatment (20 lines total) from the populations established by (Roberts et al., 2020) based upon their history of population bottlenecking and health, but not due to their resistance or development time. We continued selection of these lines for >36 generations past their initial 14, so that each line had a cumulative selection time of >50 generations (corresponding to ~4 years). Note that differences in generation time compounded over the years so the generation numbers of selection lines at the second (final) set of assays varied up to 10 generations.

To continue selection, populations were reared under the same conditions as in (Roberts et al., 2020): 1000mL straight-side wide-mouth Nalgene pots (ThermoFisher Scientific, U.K.) with 200g of their appropriate food medium in separate virus and control incubators set at 27±2 °C and 35±5% humidity, with 16:8hr light: dark cycles. Each generation (~1 month), we recorded the day of first adult emergence for each selection line, cleared the pot of all adult moths 2 days later (to prevent selection for early emergence), and then moved 50 newly emerged adult moths onto a new pot of the appropriate food medium 2 days after that to establish the next generation.

### Resistance and development time assays

For each selection line, we measured resistance (proportion infected) and development time (days until pupation and mass at pupation) on both a common garden (standard food: 0% methyl cellulose) and virus-free home (CLF or CHF) environment at both the early and final time point (Boots, 2011). To prepare for assay, selection was relaxed, and populations were spilt by approximate next emergence date into 5 batches with 1 selection line from each of the 4 treatments. Once each line in the batch had enough adult moths, 50 adults from each line in the batch were moved onto new pots containing virus-free food of the appropriate quality for each set of assays. For common garden assays, all treatments were moved onto standard food. For home environment assays, VLF and CLF lines were moved onto virus-free low-quality (CLF) food and VHF and CHF lines were moved onto virus-free high-quality (CHF) food. This ‘relaxation’ step prevent maternal effects from confounding our assays (Boots and Roberts, 2012). When adult moths emerged in parental pot, 80 adult moths were moved onto a new pot of the same food type to set up an assay pot. 11 days after setting up this pot, third instar larvae were collected for resistance and development time assays.

For resistance assays, 200 third instar larvae from each selection line were exposed to four 10-fold dilutions of virus (50 larvae dosed per dilution) corresponding to LD0-LD60. The 50 third instar larvae for each selection line x dose x assay combination were collected in separate petri dishes and then starved under a damp paper towel for 1 hour. Tiny droplets of virus solutions containing the appropriate dilution, 2% sucrose, and 0.2% Coomassie Brilliant Blue R-250 dye (ThermoFisher Scientific, U.S.A.) were placed into each dish using a syringe. The sucrose entices the larvae to consume the virus and the dye allows for confirmation that larvae have ingested half their body length of solution. These larvae were considered successfully exposed and used to fill 25-cell compartmentalized square petri dishes (ThermoFisher Scientific, U.S.A.) filled with the appropriate food type (standard for common garden assays, high or low-quality for home environment assays) with 1 grid plate per selection line x dose x assay (home or common garden) combination and 1 exposed larva per cell. Assay grids were then placed into a single incubator at the same conditions that populations were reared and allowed to develop. After 21 days, grids were frozen and destructively sampled to measure the proportion of larvae infected and uninfected. Infected larvae were distinguishable as a successful PiGV infection turns the larvae opaque, chalky white, while non-infected larvae will continue their life history as normal to pupate.

For development time assays, 100 3^rd^ instar larvae from each selection line were placed individually into four 25-cell compartmentalized square petri dishes containing either common garden or home food with 1 larva per cell and 2 grids for each of the assay environments. Grids were then moved to a single incubator at the same conditions that populations were reared in and checked every other day. Day of pupation was recorded for each larva and then each pupa was weighed 2 days later. Mass at pupation was then divided by days to pupation to calculate growth rate.

### Statistical analysis

Analyses were conducted using a generalized linear mixed modeling approach in R (“R version 4.1.2 (2021-11-01)”) (R Core Team 2021) using packages ‘lme4’(Bates et al. 2015) and ‘glmmTMB’ (Brooks et al., 2017) to build models; ‘DHARMa’ (Hartig and Lohse, 2021) to check residuals; ‘afex’ (Singmann et al., 2019) and ‘car’ (Fox and Weisberg, 2019) to determine significant model terms; ‘emmeans’ (Lenth, 2019) to extract effects; ‘tidyverse’ (Wickham et al., 2019) to manipulate data; and ‘ggplot2’, ‘ggforce’, and ‘patchwork’ to plot results (Pedersen, 2020; Pedersen and RStudio, 2021; Wickham, 2009). For all response variables, we determine error structures for models iteratively by testing fitted model residuals with ‘DHARMa’ and then adjusting error structures to best match model assumptions. We corrected residual distributions by sequentially testing models with observation level random effects (Harrison, 2014), negative binomial distributions, then zero-inflated negative binomial or quasi-Poisson distributions as needed. Once model fits were satisfactory, we tested for significance of predictor terms. Models of resistance used a binomial error structure to account for the binary outcome of ‘Infected/uninfected’. Growth rate models used gaussian, Poisson, negative binomial, or generalized Poisson error structures as required (See Supplementary model tables). Annotated R code and model output tables are attached in the supplement (note that models are named (M1-22) and that the model name that estimates and p-values are drawn from are included alongside these numbers).

For the first part of our analysis, we re-analyze the early time point (generation 14) data from (Roberts et al., 2020) for our subset of populations to see if the lineages we selected varied in their susceptibilities and growth rates in these initial assays and newly analyze the final time point (generation 50+) data to ask whether treatment affects susceptibility and growth rate after further generations of selection. For each of these time points, we separately analyze common garden assay and home food assay data sets to see if differences in a population’s conditions (virus/control and high-quality/low-quality food) led to evolved differences in resistance or growth rate. For each response variable for each data set, we ran two models to test the effects of treatment on our response variables. The first model included treatment as an interaction between ‘evolution resource environment’ and ‘virus exposed/control’ and the second model included ‘treatment’ directly. This allowed us to explore the interacting effects of our two experimental manipulations as well see the differences between individual treatments. All models include a resistance (‘proportion infected’) or life history (‘growth rate’) metric as the response variable with ‘replicate line’ nested under ‘treatment’ as random effects to account for our experimental structure. For final time point data, we include an additional random effect of ‘batch’ to account for the batched assay structure. For resistance models, ‘dose’ was included in the model as a fixed effect.

For the second part of our analysis, we ask whether the different evolution treatments lead to differences in the change in resistance or growth rate between the early and final time points. For each selection line, we calculate the change in both growth rate and resistance effect size estimates between the early and final time points. We then use the same linear modelling framework as above to determine whether ‘treatment’ informatively predicts the change in resistance or growth rate for each line in the common garden and home assay conditions between generation 14 (early) and the end of the experiment (final).

For the final part of our analysis, we test whether a given line’s growth rate is significantly predicted by its measured degree of resistance to infection, and whether this changes over time. We again analyze common garden and home assay data separately using the same linear modelling framework as above with ‘growth rate’ as the response variable and ‘susceptibility estimate’, ‘treatment’, and ‘time point’ as fixed effects with potential interactions.

## Results

### Selection Line Resistance

For the subset of lines that we further select, lines from virus selected conditions have significantly lower (estimate = −0.75, p = 0.02, Supplementary Model Tables M4) susceptibility to infection when assayed on home food at the early time point. This effect is primarily driven by the decreased susceptibility of populations from the virus high-quality (VHF) treatment (estimate = −0.75, p = 0.03, M3), as the virus low-quality (VLF) populations do not have significantly lower (estimate = −0.19, p = 0.57) susceptibility than control lines. However, this effect does not hold when lines are assayed in the common garden environment as neither ‘treatment’ (p = 0.15, M1) nor ‘virus/control’ (p = 0.95, M2) significantly affects proportion infected in these models. Across all models at the early time point, assay virus ‘dose’ significantly affects proportion infected (p < 0.001, M1-4) and neither ‘evolution food type’ (common garden: p = 0.09, M2; home: p = 0.081, M4) nor the interaction effect between ‘evolution food type’ and ‘control/virus’ (common garden: p = 0.95, M2; home: p = 0.59, M4) have significant effects. Therefore, selection from virus led to lower proportions infected at this early time point when populations were assayed in the home food environment, but not when assayed in the common garden environment (Fig 1 A-B, E-F). We did not, however, see a significant effect resource quality on evolved resistance at the early time point (Fig 1 A-B, E-F).

**Figure 1:**
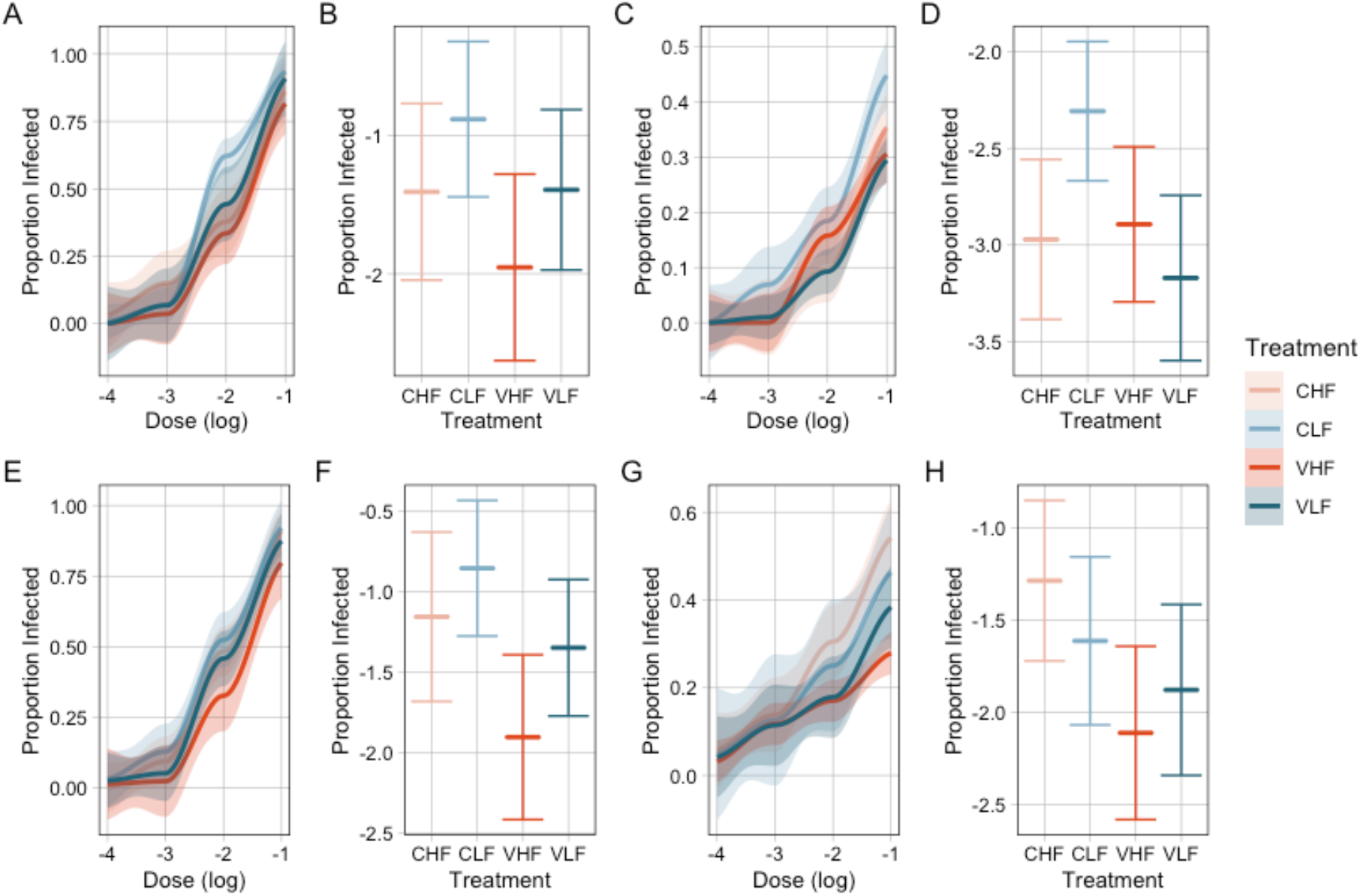
Resistance of selection lines after 14 (A-B, E-F) and 50+ (C-D, G-H) generations of section when assayed in common garden (A-D) and home food (E-H) environments. Panels A, C, E, F show smooths of the raw proportions infected at each dose, with dose log transformed on the x-axis. Panels B, D, F, G present model estimates for each treatment’s effect on proportion infected across doses from models M1, M3, M5, M7 (See Supplementary Model Tables).

After additional generations of selection, lines from virus selected conditions continue to have significantly lower susceptibility to infection in the home food assays (estimate = −0.83, p = 0.03, M8). In this case, neither ‘evolution food type’ nor the interaction between ‘evolution food type’ and ‘virus/control’ is significant (evolution food type: p = 0.84, M8; interaction: p = 0.227, M8) and the effect is driven by both virus high-quality (estimate = −0.83, p = 0.01) and virus low-quality (estimate = −0.59, p = 0.06) selected treatments, though only the virus high-quality treatment is statistically significant on its own. However, in the common garden assays, there is a strong interaction effect between ‘evolution food type’ and ‘virus/control’ (p = .009, M6) so that only control low selected populations significantly differ in their susceptibility to virus infection (estimate = 0.66, p = 0.003, M5). In these common garden assays, virus low-quality and virus high-quality lines are no more resistant than the control high-quality lines. Across all models at the final time point, the virus dose assayed at significantly affects proportion infected (p < 0.001, M5-8). Therefore, further selection from virus continues to result in lower susceptibility to infection when assayed on home food conditions, but not common garden conditions (Fig 1 C-D, G-H). After additional evolution, however, this result is not as independently driven by the lower susceptibility of virus high-quality lines, as virus low-quality lines have gotten closer to the virus high-quality lines in resistance, though still not ‘caught up’. This suggests that the context dependent resistance seen at the earlier time point is not a transient evolutionary strategy and that the dual stressor effect of low-quality resources preventing the evolution of resistance diminishes over time.

Finally, ‘treatment’ does not predict a selection line’s change in susceptibility between the early and final time points in either the common garden (p = 0.5) or home (p = 0.5) assays (Figure S1).

### Selection Line Life Histories

For the subset of lines that we further evolve, lines from virus selected conditions do not have differences in growth rate when assayed in common garden (p = 0.4, M11) or home quality food (p = 0.4, M13) at the early time point. There are no differences between treatments in the common garden assays (p = 0.8, M12), but treatments do significantly vary in the home assays (p = 0.007, M14). This is driven by lines evolved and therefore assayed on low-quality food developing more slowly (estimate = −0.75, p < 0.001, M13). Therefore, the only factor affecting growth rate at the early time point is whether the line is being assayed on low-quality or high-quality food, and the evolved resistance of virus high-quality lines in the home assays (Fig 1E-F) does not seem to come at a treatment-level cost of slower growth rate (Fig 2A).

**Figure 2:**
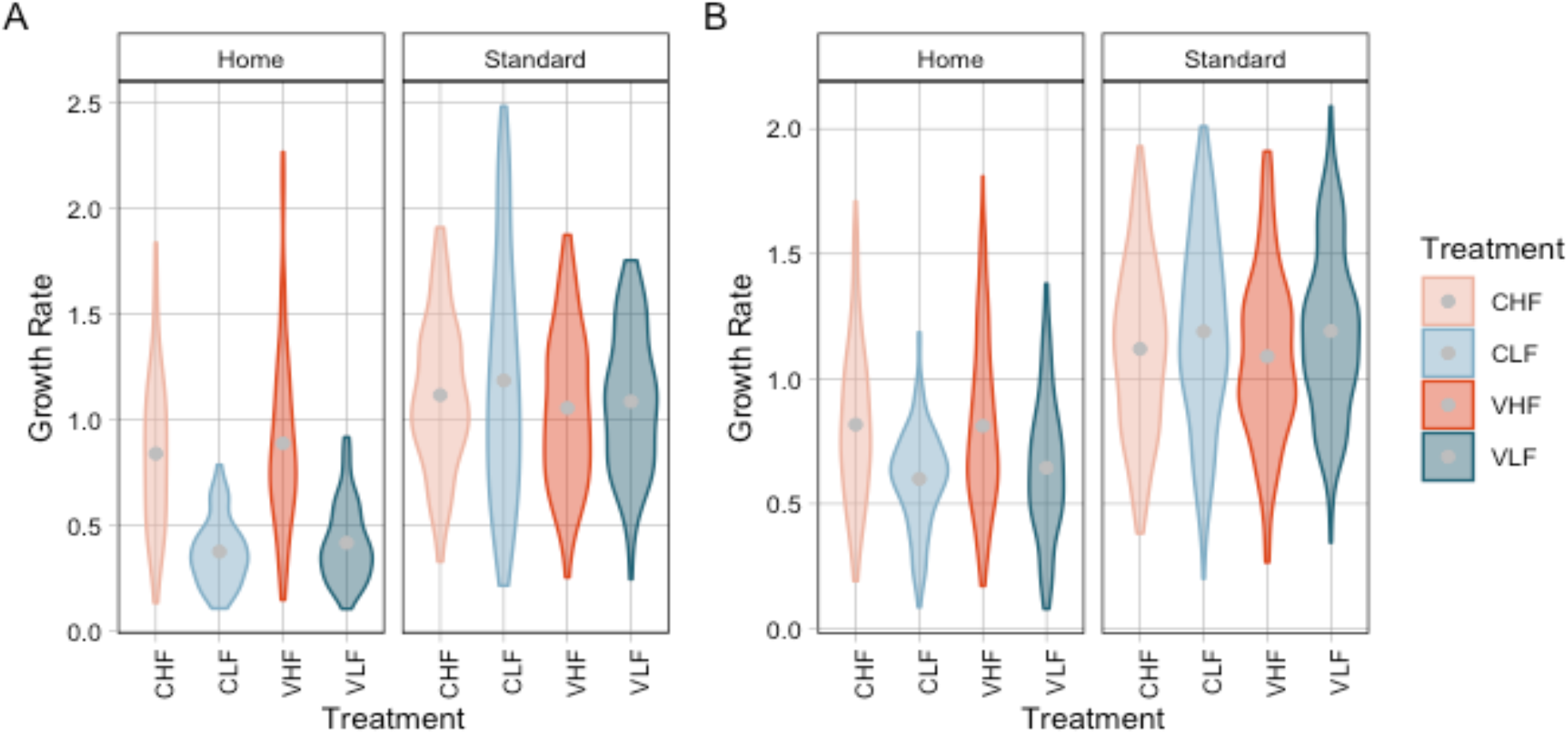
Growth rates for lines at early (A) and final (B) time points in standard and home food assays. Growth rate is the mass at pupation divided by the days until pupation. Violins represent raw growth rate data separated by treatment, assay condition, and assay time point. Gray dots represent means.

After additional generations of selection, lines from virus selected conditions continue to show no differences in growth rate when assayed in common garden (p = 0.89, M15) or home quality food (p = 0.77, M17) at the final time point. There also continue to be no differences between treatments in the common garden assays (p = 0.45, M16), and treatment is no longer significant in the home assays at the final time point (p = 0.179, M18). However, lines evolved and therefore assayed on low-quality food still develop more significantly more slowly when considered together (estimate = −0.27, p = 0.03, M17). Therefore, growth rate is still affected by whether the line is being assayed on low-quality or high-quality food at the final time point, and the evolved resistance of VHF lines, and now VLF lines, in the home assays (Fig 1G-H) continues to not be associated with a treatment-level cost of slower growth rate (Fig 2B).

Finally, treatment does not affect the change in growth rate between early and final time points in common garden assays (p = 0.8, M20, Fig S3C-D). In home assays, treatment does not have an overall significant effect on growth rate (p = 0.13, M19), but CLF lines have a borderline significant increase in growth rate (estimate = 0.52, p = 0.059, M19, Fig S3A-B).

### Correlations between Resistance and Growth Rate

In common garden assays, neither susceptibility (p = 0.15), treatment (p = 0.34), time point (p = 0.98), nor any interaction term therewithin have significant relationships with growth rate (M22, Fig3A). The sole significant single term is the interaction between susceptibility and the VLF treatment where the relationship between growth rate and susceptibility is significantly positive (estimate = 0.0005, p = 0.03) so that the fastest growers are the most susceptible.

In home assays, however, susceptibility (p = 0.005), treatment (p = 0.007), time point (p = 0.037), susceptibility: treatment (p < 0.001), susceptibility: time point (p = 0.02), and susceptibility: treatment: time point (p = 0.038) all have significant effects on growth rate (M21, Fig3B). The only non-significant model term is the interaction between treatment and time point (p = 0.11). Notably, the relationship between susceptibility and growth rate becomes significantly more negative at the final time point (estimate = −0.005, p = 0.02). From Figure 3B, we can see that this is largely because low-quality food lines shift from a positive relationship between growth rate and susceptibility in the early assays where the fastest growers are the most susceptible to a negative relationship in the final assays where the fastest growers are the most resistant. At the same time, VHF lines shift from a negative relationship between susceptibility and growth rate (fastest growers are most resistant) to a positive one (fastest growers are least resistant). We can also see that, for the same growth rate, virus selected lines from both low-quality and high-quality backgrounds are less susceptible than their control counterparts at the early time point. At the final time point this effect holds for VHF and CHF lines, but an unusually fast-growing, high-resistance CLF 5.1 means that the trend reverses for the VLF and CLF lines. Therefore, there are within-treatment level trade-offs between resistance and development time in the VLF (but not VHF) lines in the early assays, but these not only disappear, but reverse, after additional evolution.

**Figure 3:**
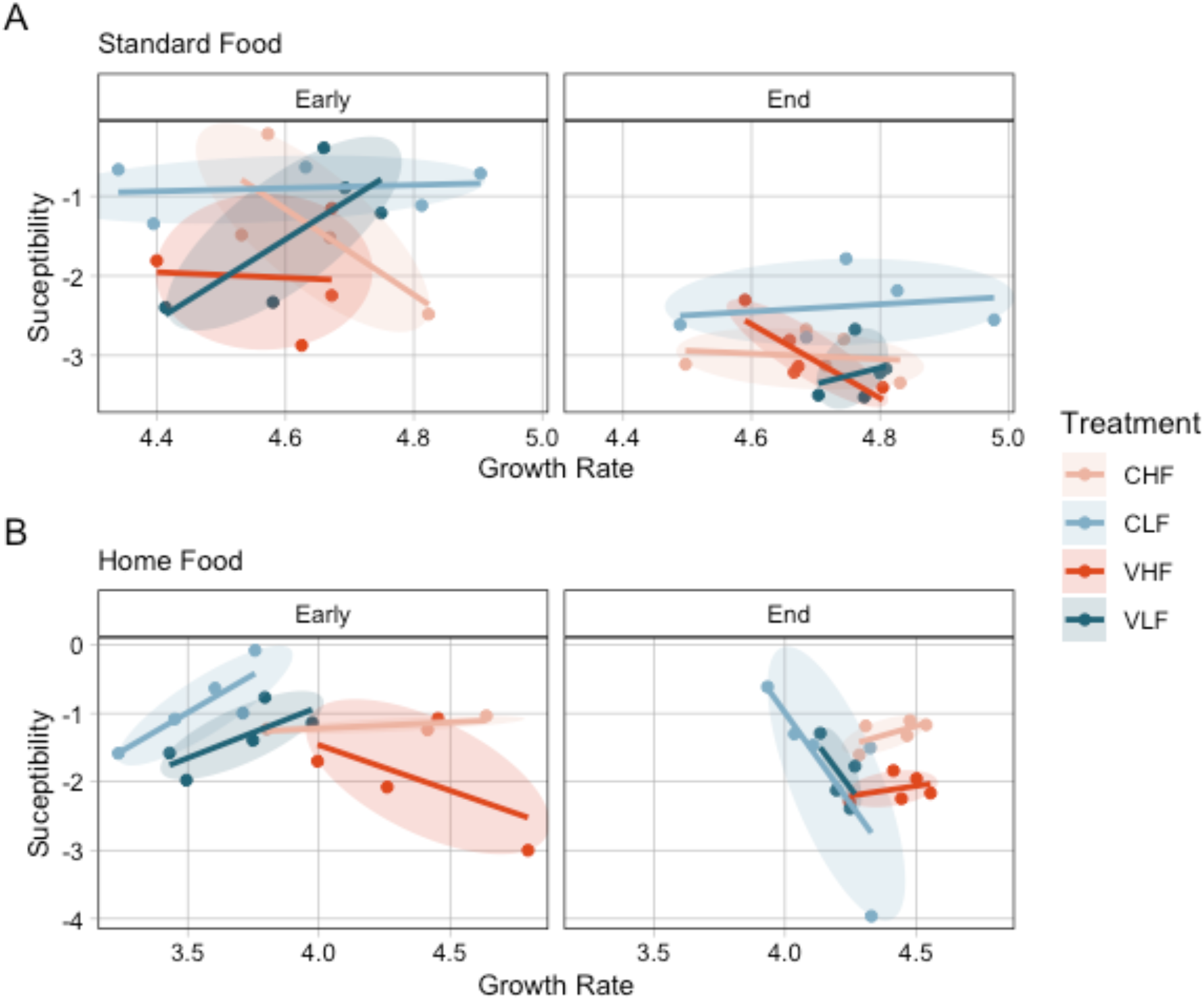
Correlations between growth rate and susceptibility in (A) common garden and (B) home environment assays at early and final time point.

## Discussion

One key result is that virus-selected lines show evidence of evolved resistance when assayed on their home food quality environment (the one in which selection occurred), but not in the common garden standard food environment. This is found at both the earlier and final time points, pointing towards a consistent effect of context-dependent resistance where selection lines’ resistance mechanisms depend on their assay environment.

Additionally, while the resistance of virus-selected lines in the home environment at the earlier time point is primarily driven by lines that were selected in the high-quality environment, after 36+ additional generations of selection, lines selected for resistance in the low-quality environment partially caught up to lines selected in the high-quality environment. Therefore, the dual evolutionary stressors of low resource quality and exposure to virus slows, rather than completely restricts, the evolution of resistance to infection. This suggests that management strategies aiming to control resistance evolution through imposing dual evolutionary stressors may only temporarily impede resistance.

Another key result is that the treatment-level pattern of costless resistance in the home environment observed in the early time point (Roberts et al., 2020) does not disappear after further generations of selection and increased resistance in the home environment. This is in contrast to previous results from the *Plodia interpunctella* and PiGV system where resistance has shown to be associated with slower growth rates (Bartlett et al., 2018; Boots, 2011; Boots and Begon, 1995, 1993). It also doesn’t follow general trends where trade-offs are more likely to be observed in longer selection experiments (Bono et al., 2017), though these trends are from microbial experiments where the number of generations is much higher. This result would suggest that the lack of detected costs was not caused by a transient effect where initial costless resistance alleles can be fixed early on, but only costly alleles remain as the population approaches the Pareto front (Li et al., 2019; Shoval et al., 2012; Visher et al., 2021). However, it is alternatively possible that our selection lines’ relatively small, bottlenecked effective population sizes mean that drift is preventing them from reaching the Pareto front and accessing costly resistance strategies (White et al., 2021). It is also possible that, since we did not sequentially increase the concentration of virus that populations were exposed to, virus-selected lines evolved ‘enough’ resistance with costless strategies and did not explore more costly resistance strategies. Still, our result that treatment-level costs to resistance don’t increase after further selection suggests that costless resistance might be a somewhat stable potential evolutionary outcome in our system and may not always be replaced by costly higher resistance strategies.

Finally, when we look at the correlations between resistance and development time within each treatment, we see that these traits can be either negatively or positively correlated depending on the time point and assay conditions. At the early time point, the selection lines evolved in the low-quality environments had positive correlations between growth rate and susceptibility in the home environment, indicating that there was a cost to resistance where the most resistant lines were the slowest developers. By the end of the experiment, however, the selection lines evolved in the low-quality environments had negative correlations between growth rate and susceptibility in the home environment, indicating costless resistance (at least in the phenotypes measured). This suggests that earlier costs to resistance were either compensated or that the low-quality selected lines started exploring different resistance mechanisms with different cost structures.

We do know that there are multiple resistance mechanisms in the *Plodia interpunctella* and PiGV system that differ in their relationships with development time (Bartlett et al., 2020; Boots and Begon, 1993). Some resistance mechanisms that involve processes like reallocating resources to increase immune investment or decreasing gut permeability come at the cost of slower development time, but faster growth rates themselves can increase resistance by shortening the time window for infection (Bartlett et al., 2020; Boots and Begon, 1993). This is because infection must establish before the gut epithelia is shed during larval molting, so shorter intervals between molts can confer higher resistance (Bartlett et al., 2020; Engelhard and Volkman, 1995; Hochberg et al., 1992). Developmental resistance mechanisms could be responsible for the negative correlations between growth rate and susceptibility in the home environment for the low-quality selected lines at the end of the experiment. Because these lines are assayed on low-quality food in home assays, they have slower growth rates that might challenge to their ability to effectively resist infection. The context dependency of evolved resistance prevents us from making good comparisons across common garden and home assays, but it is notable that the correlation between susceptibility and growth rate is significantly positive for VLF in the common garden at the end time point. Growing on standard quality food moves them higher on the growth rate axis and therefore could move them out of the region where infection interval length is very important. These results suggests that the resistance of the low-quality food selected lines in the home environment assays both quantitatively and qualitatively changes between the early and final time points.

As a whole, our results emphasize the influences of time scale, resources, and context dependency on trade-offs between life-history traits. We show that the nature and, in particular, the costs of resistance in this long-term experiment (Roberts et al., 2020) differ from previous results in the system that found consistent costs to resistance with development time (Bartlett et al., 2018; Boots, 2011; Boots and Begon, 1995, 1993). This could be explained by there being differences in the starting populations such that different resistance conferring variants existed in the standing genetic variation of the ancestral population for each experiment. Furthermore, even if the starting variation was identical, different sampling of early adaptive alleles could have led towards different historically contingent evolutionary trajectories. Finally, it does seem that selected lines were able to evolve qualitatively and quantitatively different resistance when given a longer time scale of selection. Thus, it is clear that trade-offs to resistance in our experiment are historically contingent and depended on evolutionary conditions and the time scale of selection. Therefore, it is also clear that trade-offs do not consistently evolve even over extended timescales and that more work is therefore needed to better understand their conditionality to better predict the evolution of resistance.

## Supporting information

Supplemental Model Tables

## Supplementary Figures

**Figure S1:**
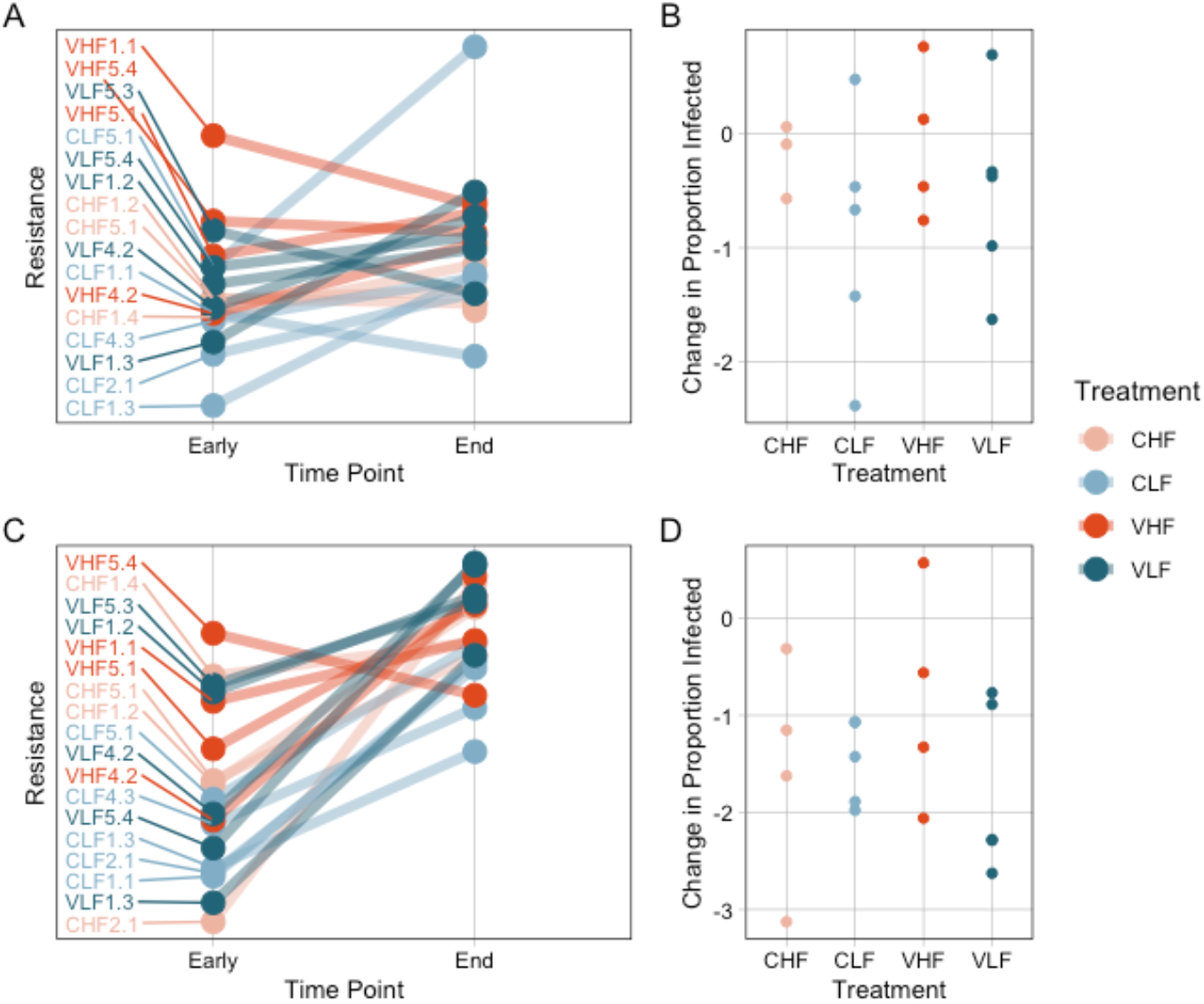
Change in resistance between Early and Final time points in Home (A-B) and Common Garden (C-D) assays. Panels A, C link resistance estimates for each line across the Early and Final time points. Panels B, D present the delta resistance values for each treatment.

**Figure S2:**
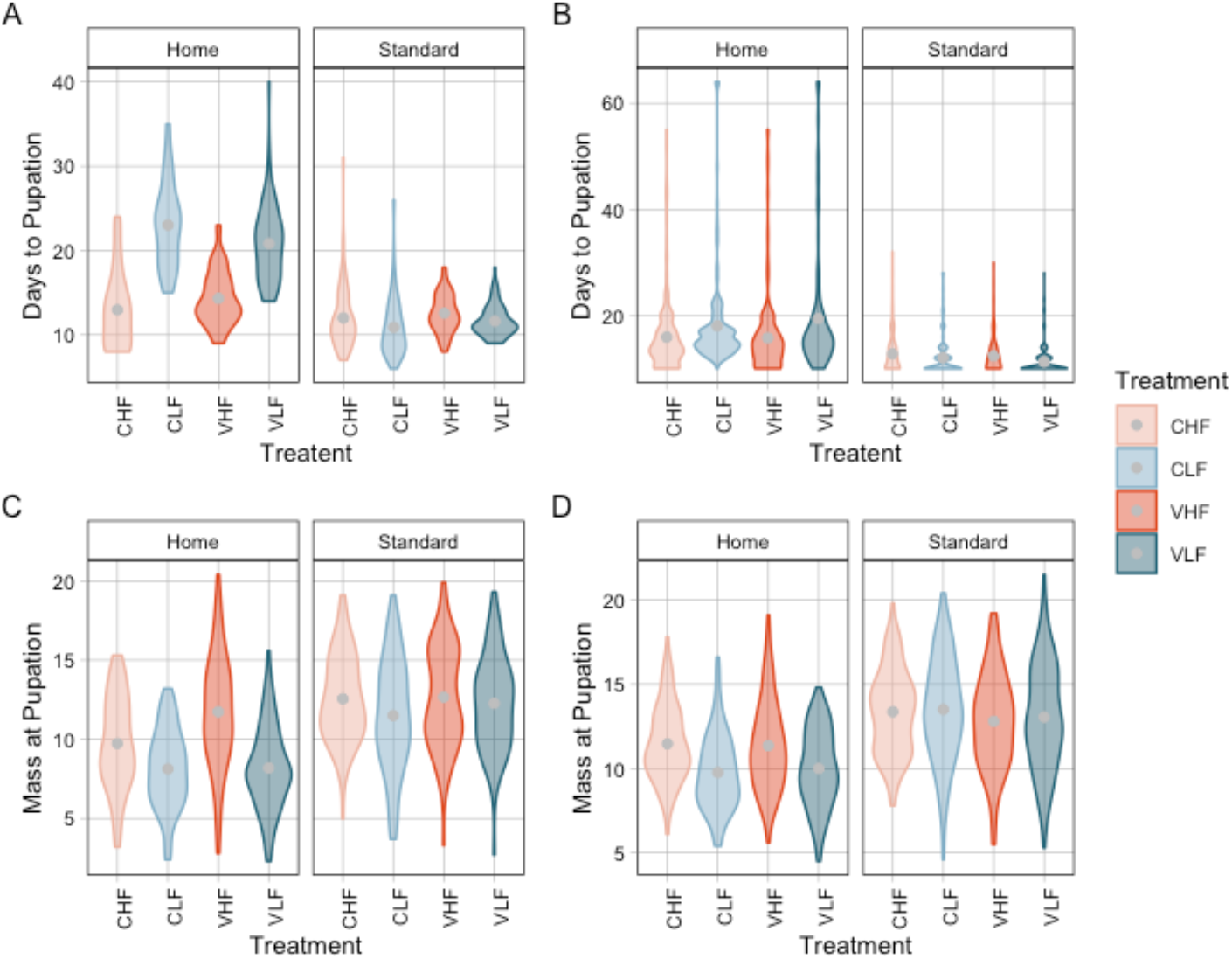
Days until pupation (A-B) and mass at pupation (C-D) for the different treatments in Home and Standard (Common Garden) assays at Early (A, C) and Final (B, D) time points.

**Figure S3:**
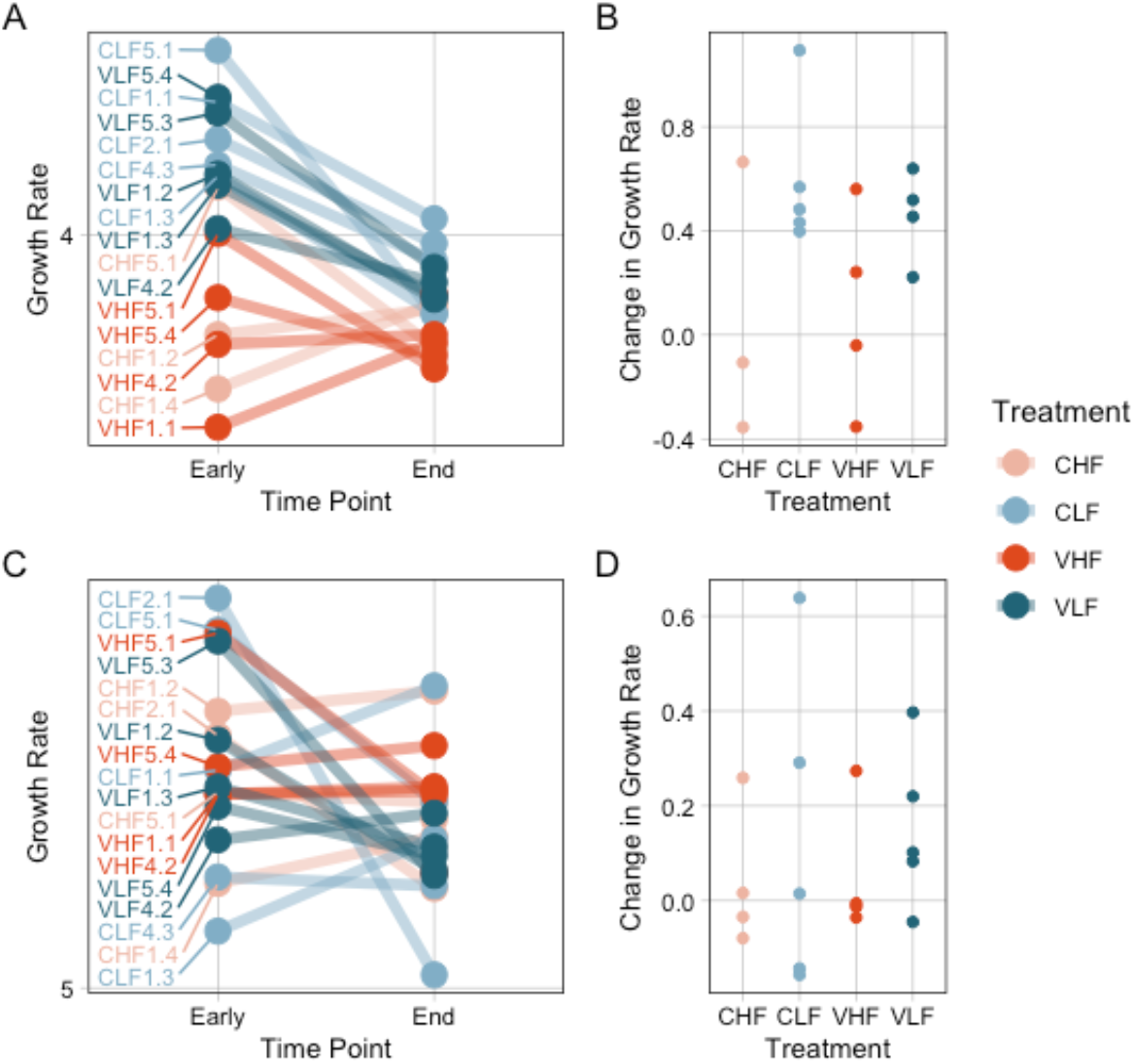
Change in growth rate between Early and Final time points in Home (A-B) and Common Garden (C-D) assays. Panels A, C link growth rate estimates for each line across the Early and Final time points. Panels B, D present the delta growth rate values for each treatment.

